# Exceeding 80% efficiency of single-bead encapsulation in microdroplets through hydrogel coating-assisted close-packed ordering

**DOI:** 10.1101/2023.02.08.527643

**Authors:** Long Chen, Yi Zhao, Jie Li, Chenwei Xiong, Yi Xu, Chengren Tang, Rong Zhang, Jingwei Zhang, Xianqiang Mi, Yifan Liu

**Author notes:** These authors contributed equally to this work (Long Chen, Yi Zhao).

## Abstract

High-efficiency encapsulation of single microbeads in microdroplets is essential for droplet-based high-throughput analysis such as single-cell genomics and digital immunoassays. However, the demand has been hindered by the Poisson statistics of beads arbitrarily distributed in the droplet partitions. Although techniques such as inertial ordering have been proven useful to improve bead loading efficiency, a general method that requires no advanced microfluidics and owns compatibility with diverse bead types is still highly desired. In this paper, we present hydrogel coating-assisted close-packed ordering, a simple strategy that improves the bead loading efficiency to over 80%. In the strategy, the raw beads are coated with a thin layer of hydrogel to become slightly compressible and lubricious so that they can be close-packed in a microfluidic device and loaded into droplets in a synchronized manner. We first show that the thin hydrogel coating can be realized conveniently through either jetting microfluidics or vortex emulsification. When loading single 30-μm polystyrene beads, we experimentally determined an overall efficiency of 81% with the proposed hydrogel coating strategy. Of note, the strategy is not sensitive to the selection of raw beads and can tolerate their polydispersity. Using the strategy, we achieve a cell capture rate of 68.8% when co-encapsulating HEK293T cells and polydispersed barcoded beads for single-cell transcriptomics. Further sequencing results verify that the reversible hydrogel coating does not affect the RNA capture behavior of the encapsulated barcoded beads. Given its convenience and broad compatibility, we anticipate that our strategy can be applied to various droplet-based high-throughput assays to drastically improve their efficiency.

## Introduction

In recent years, droplet microfluidics has been extensively involved in high-throughput biotechnologies and relevant biomedical research^1-4^. A variety of impactful techniques have been developed based on the droplets, such as single-cell sequencing^5-8^ and digital molecular assays^9^. All of these techniques rely on droplets because they can serve as isolated, minute, and manipulable reactors ideal for massive independent biochemical assays^10, 11^. The droplets can be formed at kHz, yielding millions of compartmentalized picolitre reactors to house single cells and molecules. Moreover, the development of microfluidic toolboxes has allowed droplets to be manipulated through techniques such as droplet merging^12-14^, dividing^15, 16^ and sorting^17-19^, and liquid injection^20^. This has enabled a wealth of complex assays to be built on droplet microfluidics, such as single-cell genomics^21^, high-throughput screening^22^, and digital immunoassays^23^.

Many droplet-based techniques require introducing functional microbeads into the droplets^17, 19, 24, 25^. The beads serve as a vehicle to carry essential biomolecules and probes, such as oligonucleotides and antibodies, to enable customized reactions in the droplets. For instance, Macosko et al. used polystyrene microbeads with immobilized DNA oligonucleotides to capture and barcode the transcriptome of a single cell in a microdroplet, thereby realizing high-throughput single-cell transcriptome sequencing^26^. Cohen et al. introduced antibody-coated microbeads to droplets to realize droplet digital enzyme-linked immunosorbent assays (ddELISA), which allows the analysis of single protein molecules^27^. In general, microbeads used in droplet-based assays can be categorized into two types based on their mechanical property: one is soft and elastic, typically made of hydrogels; the other is hard and rigid, which is composed of glass, plastic or ceramic materials. A major advantage of hydrogel beads is that they can be close-packed in a microfluidic channel to achieve an encapsulation process synchronized to droplet generation, thereby ensuring a high overall reaction efficiency (% droplets containing an effective reaction)^28-30^. On the other hand, hard beads are advantageous in that they are compatible with both aqueous and organic solvents, which is particularly crucial when DNA oligos are to be immobilized on the beads as oligonucleotides can be directly synthesized on the surfaces via phosphoramidite chemistry. Moreover, the vast variety of solid materials can introduce more desired properties such as being magnetic^19^ and fluorescently encoded^31^. Benefiting from these unique features, the hard microbeads have been a major choice for many representative droplet-based assays^26, 27, 31^. However, there is an intractable issue with the hard beads that loading a single hard bead into a single droplet relies on limited dilution and is therefore subjected to the Poisson distribution^5, 32^. This leads to less than 10% of the droplets containing a bead and thus a low reaction efficiency. Efforts have been paid to order the beads using principles such as inertial ordering^33, 34^, and hydrodynamic focusing^35^. However, such methods may require extreme flow conditions^36^, sophisticated fluidic designs^33, 34^, and sometimes additional sheath flows^35^. These extra requirements limit their potential to be practically applied, especially in a commercialization-aimed setting where simplicity and robustness are critical. Therefore, there is an urgent need to develop a convenient and user-friendly strategy to order these microbeads in microfluidics for higher-efficiency droplet-based assays, particularly a strategy that is directly compatible with existing droplet microfluidic systems.

Here, we report a simple, robust, and scalable approach to realize super-Poisson loading of rigid microbeads into droplets. In this approach, we coat a thin layer of hydrogel around a hard microbead, rendering the beads appreciably soft, lubricous and compressible. Therefore, a key feature of the hydrogel-coated rigid beads (HRBs) is that they can be close-packed in a microfluidic channel to beat the Poisson distribution just as the complete hydrogel bead does. By squeezing and close-packing, the beads can be loaded to the droplets with a regular interval in synchronization with the droplet formation; this results in a super-Poisson distribution in that the majority of droplets contain a single bead, Fig. 1. Benefiting from this, the reaction efficiency is increased dramatically. Considering the yield quality and scalability, we developed two approaches to fabricate the HRBs. The first approach relies on jetting microfluidics where the existence of a microbead triggers the breaking of a thin liquid jet due to the Rayleigh-Plateau instability. This leads to HRBs with a high monodispersity. The second approach relies on particle-templated vortex emulsification which yields slightly polydispersed HRBs but features a higher manufacturing capacity and scalability and does not require microfluidics. Experimentally, we obtained a loading efficiency (ratio of droplets containing a single bead) of 81.0% when using microfluidically manufactured HRBs and a simple flow-focusing device with no complicated fluidic designs, over 8-fold higher than that of the limited dilution-based loading of the bare rigid beads. Even using non-uniform beads as templates (e.g. Drop-seq beads^26^), a high loading efficiency of 68.8% was achieved, which outperforms previously reported internal flow-based approaches^33, 36^. Of note, the hydrogel coating does not compromise the functionality of the beads. Once encapsulated in the droplets, the coating can be melted by gentle heating, exposing the immobilized biomolecules. High-throughput sequencing reveals that HRBs prepared from Drop-seq beads^26^ are able to capture RNA transcripts released from single cells. Given their convenient fabrication, outstanding performance, and compatibility with conventional droplet microfluidics, we anticipate that HRBs can be applied to various droplet-based high-throughput techniques to drastically increase their reaction efficiency while keeping their original design of microfluidics.

**Figure 1.**
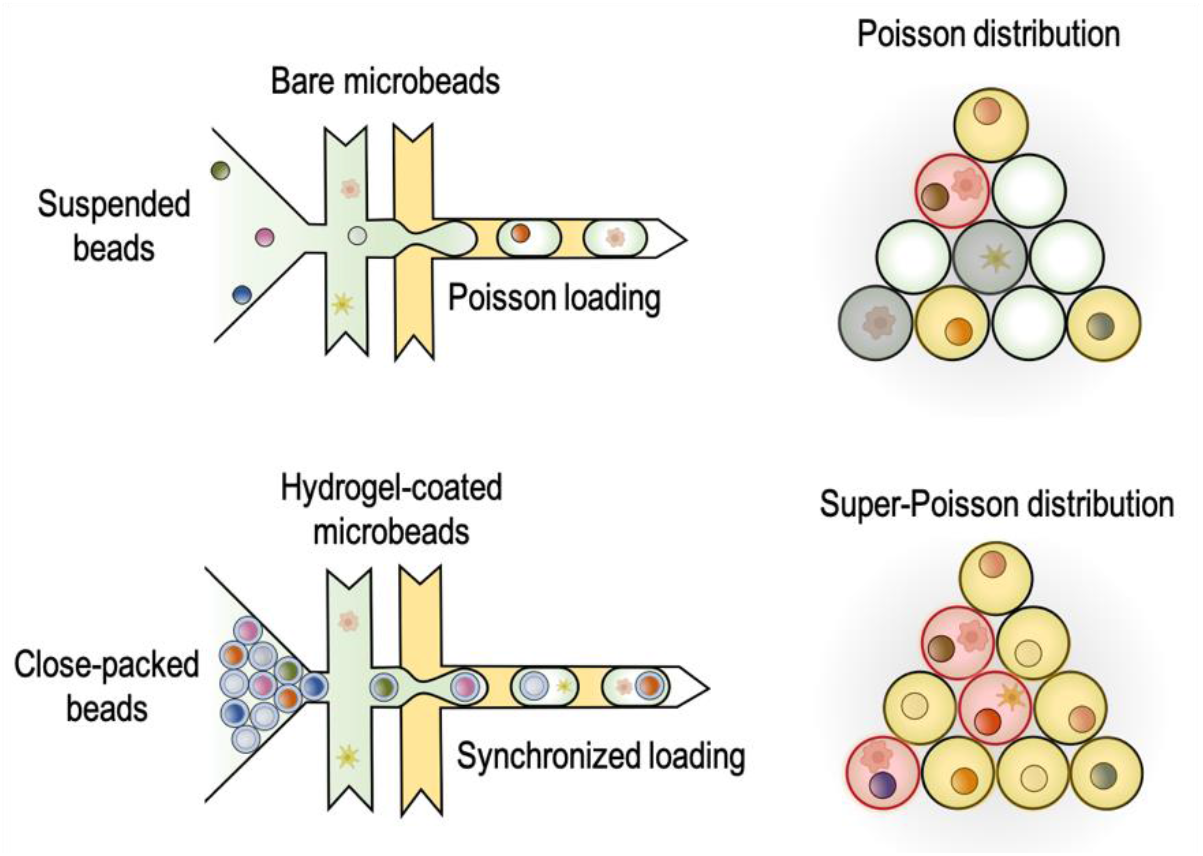
Conceptual illustration of high-efficiency bead loading achieved by hydrogel coating-assisted close-packed ordering. Limited dilution-based bead loading is subjected to Poisson distribution and is thus low-efficient because most droplets are empty. This results in severe waste of reagents and precious biological samples (for example cells in the illustration). The presented hydrogel-coated beads can be close-packed in a microfluidic channel to be loaded into droplets in a synchronized manner, thus achieving high-efficiency super-Poisson loading.

## Materials and methods

### Device Fabrication

The fabrication of the microfluidic device was carried out using standard polydimethylsiloxane (PDMS)-based soft lithography^5^. The layout of microfluidic devices was sketched using AutoCAD and printed as dark-field plastic film masks. To fabricate the silicon mold, a negative photoresist SU-8 3025 (MicroChem) was spin-coated on a 3-inch silicon wafer to the desired thickness. After a pre-bake, the wafer was covered with the mask and exposed under a 120-mW UV lamp (Thorlabs, M365L2). Then, the wafer underwent a post-exposure bake and was developed in a SU-8 developer solution (MicroChem). The baking durations and exposure dosage followed the manufacturer’s instructions. The fabricated mold was cleaned with isopropanol and ethanol and was blow-dried using a nitrogen gun. Next, PDMS precursor (SYLGARD 184, Dow Corning) was mixed with its curing agent at a ratio of 10:1 (w:w) and degassed in a vacuum chamber. The prepared PDMS was poured over the mold, followed by a curing process at 60 °C for overnight. The cured PDMS slab was peeled off from the mold, and inlet/outlet ports were created using a 0.7 mm custom-made hole puncher. The slab was then bonded to a clean glass slide *via* oxygen plasma treatment. The resulting chip was baked at 100 °C for 30 min to enhance the bonding quality. Before use, the devices were treated with a commercial water repellent, Aquapel (PPG Industries), to render the channel surfaces hydrophobic.

### Manufacturing of HRBs

Two kinds of microbeads were selected to fabricate HRBs: 1) 30-μm-diameter uniform polystyrene beads and 2) non-uniform, oligonucleotide-functionalized microbeads (ChemGene, Cat#MACOSKO-2011-10) used in the Drop-seq^26^ for RNA transcripts capture and barcoding. To manufacture HRBs, the microbeads were first added into a warm agarose solution (2% wt, Sigma-Aldrich, Cat#A9414) to a final concentration of approximately 200 per μL. The bead suspension was kept on a heating block (at 70 °C) to prevent gelling and vortexed thoroughly before emulsification. To emulsify the suspension using jet microfluidics, it was transferred to a 1-mL syringe. The syringe and another 2-mL syringe containing the oil phase were further connected to a microfluidic flow-focusing device (Fig. S1) through plastic tubings (Scientific Commodities Inc., Cat#BB31695-PE/2). To conduct the experiment, the device was placed on an inverted microscope (Nikon Eclipse Ti2) equipped with a high-speed camera (Phantom Veoe-310L) for observation and the syringes were mounted on syringe pumps (New Era, NE-501). During the emulsification, the syringe and device were kept warm with a house hot air heater being set at a 20 cm distance. The flow rates were set as 2.6 mL/h and 26 mL/h for the water and oil phases, respectively, to generate a stable jet. The resulting droplets were collected in a centrifuge tube set on ice and further incubated at 4 °C for 30 min to ensure the solidification of the microgels. To emulsify using vortexing, 500 μL of the prepared bead suspension was pipetted into a 1.5-mL centrifugal tube pre-filled with 500 μL of oil. The tube was then secured on a vortexer (DLAB, MX-S) and vortexed for 10 min at 2300 rpm. The resulting emulsion was kept at 4 °C for 30 min. For both approaches, a fluorinated oil (3M HFE-7500) with 2% (w/w) amphiphilic copolymer surfactant (ThunderBio) was used as the oil phase. To break the emulsion, 2 mL of 10% (v/v) perfluoro-octanol (Sigma Aldrich, Cat#370533) in HFE-7500 was added to 1 mL of the droplet suspension and mixed by pipetting. The mixture was centrifuged at 1,000 g for 1 min and the bottom oil layer was discarded. This process was repeated once to guarantee the removal of excessive oil. Then, the remaining gel beads were washed thrice with 0.01% (v/v) Tween 20 (Bioss, Cat#C-0086) and Tris-EDTA (TE, pH 8.5) buffer, respectively. To enrich the HRBs, the gel beads were resuspended in TE buffer and centrifuged at 300 g for 1 min, during which the HRBs were spined down quicker than empty gel beads and thus settled at the very bottom. Next, the supernatant and upper layer (∼ 90%) of gel beads were discarded. This process was repeated thrice to obtain high-purity HRBs. The HRBs were further filtered twice using a 70-μm cell strainer (Falcon, Cat#352350) and a 40-μm cell strainer (Falcon, Cat#352340), respectively.

### Bead loading experiments

The prepared HRBs were first spined down (1000 rpm, 1 min) to remove excessive buffer solution, after which the dense beads were transferred to a 1-mL syringe. The HRBs were reinjected to a dual-junction flow-focusing device (Fig. S2) at a fixed flowrate of 60 μL/hr. The flowrates for the two aqueous flow inlets and the carrier oil inlet were 70 and 220 μL/hr, respectively. The aqueous phase was either a 1X phosphate-buffered saline (PBS) solution or a cell suspension solution. The Poisson loading of beads in droplets was estimated by the Poisson distribution equation, *P*(*x*) = *λ ^x^ e ^−λ^*/*x* !, where *P(x)* stands for the possibility of *x* beads in a droplet and λ describes the mean number of beads per droplet (experimentally obtained).

### Cell culture and preparation

Human embryonic kidney (HEK) 293T cells were cultured in Dulbecco’s Modified Eagle Medium (DMEM, Thermal Fisher, Cat#12491015) supplemented with 10% fetal bovine serum (FBS, Thermal Fisher, Cat#26140) and 1% Penicillin Streptomycin (Thermal Fisher, Cat#15140148) incubated in a 5% CO_2_ humidified atmosphere at 37°C. The cells were harvested by trypsinization and then filtered twice with a 40-μm cell strainer. The density of collected cells was measured by a hemocytometer (JIMBIO-FIL, China). For the bead-cell co-encapsulation experiments, the cells were spined down (1500 rpm/min, 1min) and resuspended in PBS buffer containing 0.01% wt bovine serum albumin (Sigma-Aldrich Cat#A8806) and 16% (v/v) Optiprep (Sigma-Aldrich, Cat#D1556) for density matching. For the Drop-seq library construction, the original Drop-seq procedures^26^ were followed unless otherwise specified.

### Bioinformatics

The constructed complimentary DNA (cDNA) fragments were sequenced on an MGISEQ-2000 sequencer. The obtained sequencing data were quality-checked (using FastQC) and processed to generate an expression matrix using the UMI-tools (https://github.com/CGATOxford/UMI-tools). Reads in the whitelist were aligned to the reference genome (Ensembl genome browser, homo_sapiens, GRCh38.103) using STAR Pipelines. This output was then imported into the Seurat (4.0.1) R toolkit for quality control and downstream analysis of the single-cell RNA-seq data. All functions were run with default parameters unless specified otherwise. We excluded barcode groups with less than 200 or more than 10,000 detected genes.

## Results and discussion

### Manufacturing of HRBs

We first determined that the proposed HRBs should meet three requirements: 1) the surrounding hydrogel layer should be thin, being sufficient to perform close-packed ordering but possessing negligible impacts on final droplets (e.g. size and chemical conditions); 2) the hydrogel applied and its gelling/dissolving process should be biocompatible to eliminate any influences on the biomolecules attached to the beads; 3) additionally, the manufacturing should be scalable to meet the high throughput of droplet assays. To achieve these goals, we designed two strategies to manufacture HRBs, which rely on jetting microfluidics and vortex emulsification, respectively (Fig. 2a). In the first approach, the presence of a bead in a thin jet induces a local curvature that aids to break the jet due to Rayleigh-Plateau instability^37^. This results in a bead-containing droplet having a thin outer aqueous layer. By contrast, the droplet generated under the dripping regime is much larger than the encapsulated bead (typically > 2 folds in diameter). To test the jetting approach, we first designed a microfluidic flow-focusing device (Fig. S1) that features an extra spiral loading channel to preliminarily order the beads using inertial flow^38-40^. The device was examined under various flow conditions to determine the dripping and jetting regimes (Fig. 2b). To obtain a stable jet, the flowrates of 2.6 and 26 mL/h were chosen, for the continuous oil phase (Q_o_) and the discrete aqueous phase (Q_w_), respectively (Fig. 2b, inset). An alternative approach to manufacturing the HRBs is bead-templated vortex emulsification (Fig. 2a)^41^. Although this mechanism leads to polydispersed droplets, the size distribution of the droplets can be shaped by the encapsulated beads when the aqueous layer is adequately thin. We next moved to fabricate and characterize the HRBs using the two approaches. We chose 30-µm polystyrene beads as raw beads and agarose as the hydrogel material due to its biocompatibility and mild gelling/dissolving condition. Fig. 2c depicts the generated emulsion droplets and the corresponding HRBs obtained *via* a centrifugal purification process. Overall, both approaches can yield HRBs with high purity (> 98%, Fig. S3). As expected, the jetting microfluidics yields HRBs with a narrower size distribution (52 ± 3 µm, n = 306), which is desired for high-efficiency close-packing (Fig. 2d). Moreover, we found that the confinement effect of the thining jet at the focusing junction can effectively prevent the formation of doublets (two adjunct beads being trapped in a single droplet), Fig.2e. As a result, the doublet rate of the final HRBs is only 0.1% (Fig. 2f). Although the vortex emulsification leads to non-uniform HRBs (45 ± 8 µm, n = 332) and a higher doublet ratio (5.97%), it is microfluidic-free and less time-consuming (Fig. 2g). In our experiments, ∼200,000 HRBs can be manufactured with vortexing in less than 10 min. If needed, scaling up the vortexing approach is straightforward, which simply requires a larger vortex mixer to hold multiple tubes for parallelization. Such scalability is highly desired for ultrahigh-throughput single-cell or digital diagnostic assays^42^, especially those undergoing a commercialization pipeline.

**Figure 2.**
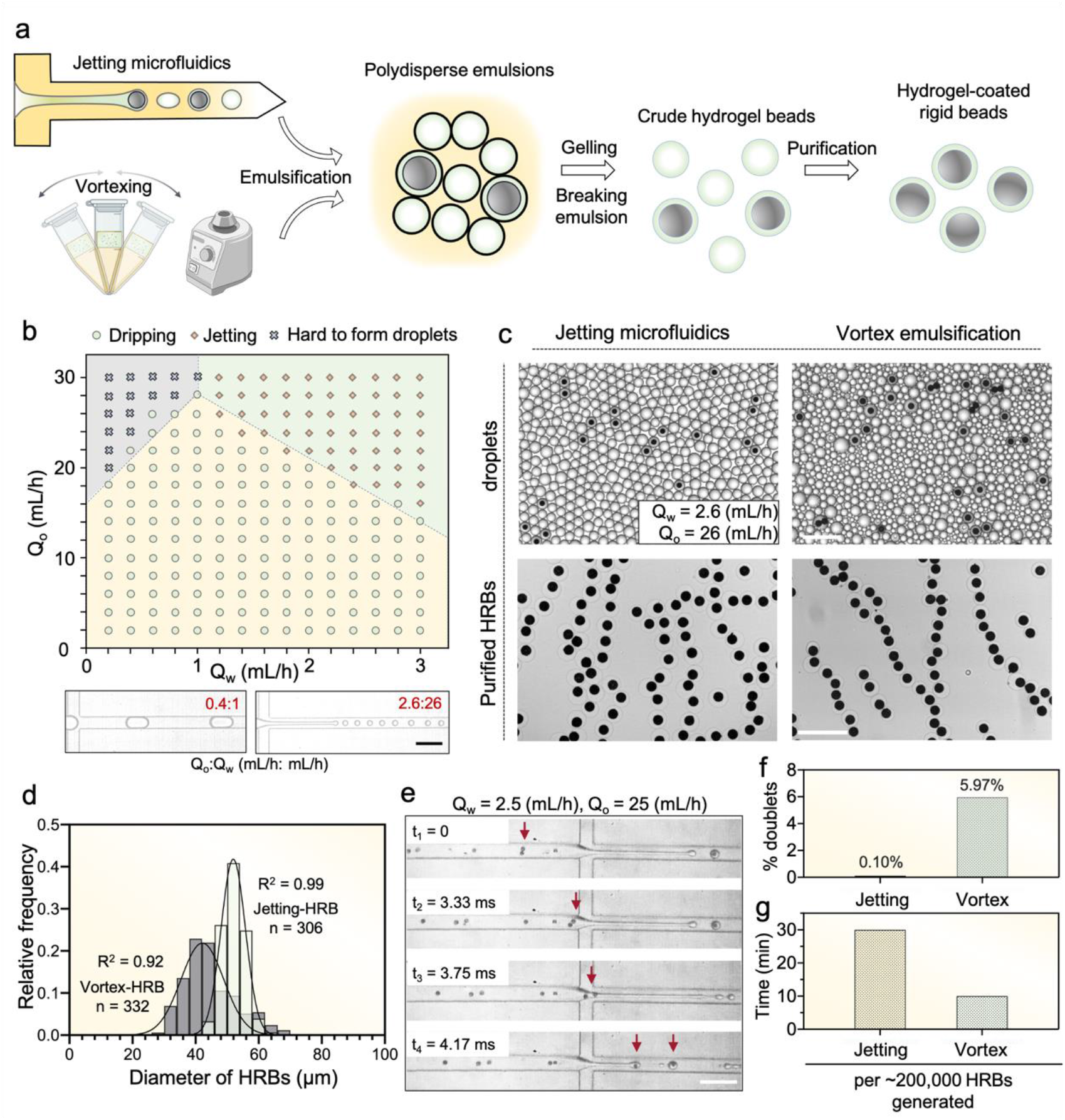
Manufacturing and characterization of HRBs. (a) The workflow of fabricating HRBs. Two alternative approaches, 1) jetting microfluidics, and 2) vortex emulsification are proposed to yield HRBs. (b) Experimentally obtained dripping-to-jetting transition map of the jetting microfluidic device. The micrographs on the bottom depict representative dripping and jetting conditions at the stated water (Q_w_) and oil (Q_o_) flowrates. (c and d) The crude droplets and final HRBs obtained from the jetting and vortexing approaches, respectively (c), and their size distribution (d). The curved lines in (d) represent corresponding Gaussian fitting results. (e) A series of micrographs showing two adjunct beads being separated by the thining jet during HRB generation.(f) Comparison of doublet rate (n = 306 for jetting and n = 332 for vortex) and consumed time between the two approaches. All scale bars: 200 µm.

### Close-packing HRBs into droplets

Having successfully manufactured the HRBs, we proceeded to test whether the hydrogel coating could lead to efficient close-packing as we proposed. To perform the experiments, we designed a microfluidic device featuring two consecutive flow-focusing junctions (Figs. 3a and S2). In this design, the close-packed beads should meet the two pinching aqueous flows at the first junction and thus be spaced at regular intervals before entering the second junction where the droplets are to be formed. In the experiments, we first observed that the HRBs were indeed close-packed in the upstream buffering channel, moved smoothly to the junction, and were encapsulated in the droplets (insets of Fig. 3a and Supporting Movie S1). We then generated the droplets using both jetting- and vortex-HRBs (Fig. 3b) and found that they are highly monodispersed, the sizes of which exhibit a typical Gaussian distribution (Fig. 3c). Based on the Gaussian fittings, we obtained that the droplet sizes are 89 ± 3 (n = 326) and 94 ± 4 (n = 313) µm for the jetting - and vortex-HRB droplets, respectively. Next, we measured the total encapsulation efficiency of both HRBs (Fig. 3d). Excitingly, we found that for the jetting-HRBs, 81.0% of the droplets contained a single bead. Meanwhile, the rate of multiple beads being encapsulated into the same droplet is only 1.5%. Such a performance presents a significant improvement over the Poisson loading performed at the same bead density (Fig. 3e). The encapsulation of vortex-HRBs exhibited a lower efficiency of 52.5% and a higher rate of multiplets, 5.3% (Fig. 2d). This is a result of the irregular loading of beads in the droplets given vortex-HRB’s variation in size. However, the overall bead encapsulation efficiency still outperforms the Poisson distribution (Fig. S4). For benchmarking, we summarized the bead encapsulation efficiency of several recent reports on single-bead encapsulation for digital assays (Fig. 3f). In terms of bead encapsulation efficiency, our work is over 6-fold higher than a recent report^27^ which solely relies on Poisson loading, and is advantageous over the reported inertial ordering-based techniques^35, 36^. This suggests that the presented strategy should be potentially interesting for high-efficiency droplet-based digital assays such as ddELISA^27^.

**Figure 3.**
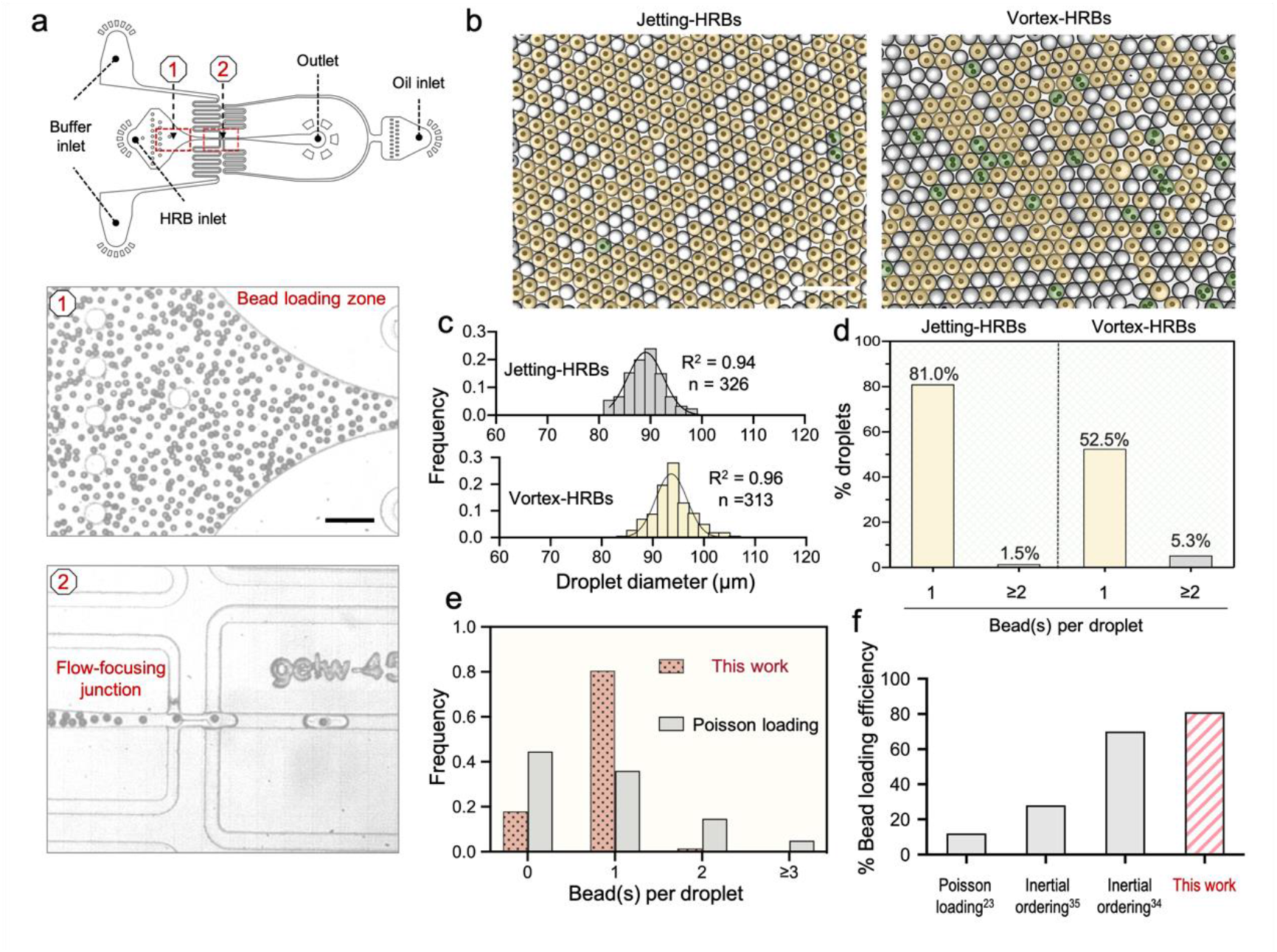
High-efficiency single-bead loading into microdroplets. (a) The layout of a flow-focusing bead loading device (top) and micrographs showing the bead loading zone and the flow-focusing junction (bottom). (b) Microdroplets generated with the jetting-HRBs and vortex-HRBs. The color shadings highlight droplets having a single bead (yellow) and multiple beads (green). (c and d) Size distribution histogram (c) and bead loading statistics (d) of the droplets. The curved lines represent the Gaussian fitting results. (e) Comparison of bead loading statistics between the Poisson loading (from calculation, λ = 0.81) and the presented strategy (using jetting-HRBs). (f) The bead loading efficiency of representative works relying on Poisson loading and inertial ordering, and the presented work.

### Bead-cell co-encapsulation for high-efficient single-cell analysis

One of the major demands for high-efficient single-bead encapsulation arises from droplet-based single-cell analysis such as single-cell RNA sequencing^26, 29, 30^. Hence, after verifying the bead encapsulation alone, we moved to test the co-encapsulation of HRBs and cells. To recapture the exact condition of a single-cell sequencing experiment, we employed the barcoding beads in the Drop-seq, a representative single-cell sequencing technique, to manufacture HRBs (Fig. 4a). The Drop-seq beads pose a severe challenge of efficient loading because they are rigid and non-uniform in size (Fig. 4b, micrographs). Interestingly, however, we found that the manufactured HRBs exhibited higher size uniformity (Fig. 4b, histogram) compared to raw Drop-seq beads. The Gaussian fitting indicated that the bare Drop-seq beads were 37.4 ± 5.4 µm (n = 82) in diameter, whereas the jetting-HRBs (fabricated out of the Drop-seq beads) were 58.5 ± 3.3 µm (n = 82). This is desired because, with a higher size uniformity, we expect the bead loading to have higher efficiency^28^. In short, we found that the hydrogel coating not only enables bead close-packing (Fig. 3) but also enhances the size uniformity if the beads are initially non-uniform, which should contribute to a higher bead loading efficiency and thus a higher rate of cells being analyzed.

**Figure 4.**
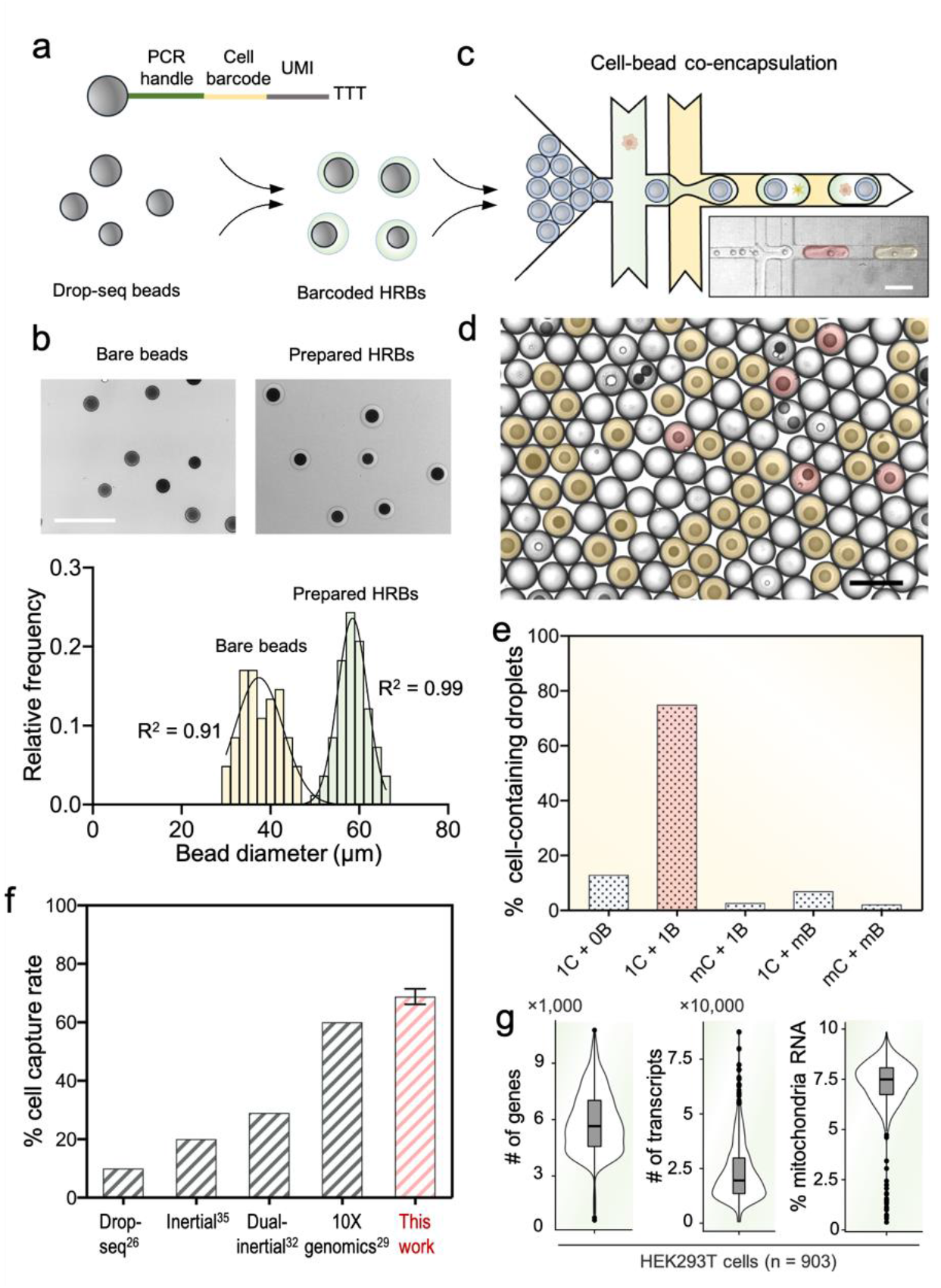
High-efficiency pairing of single Drop-seq beads and single cells using hydrogel coating-assisted close-packed ordering. (a) Schematic of a Drop-seq bead with attached oligos of the specified sequence structure (UMI: universal molecular index), which is further manufactured into an HRB. (b) Micrographs of bare Drop-seq beads and the prepared HRBs (top) and their size distribution (bottom, n = 82). The curved lines denote Gaussian fittings to each distribution profile. Schematic and a representative micrograph showing the prepared HRBs being closed-packed and loaded into droplets in a synchronized manner, while the cells are loaded from the side channels. A representative micrograph of the generated droplets. The color shadings highlight droplets having a bead only (yellow) and a bead + a cell (green). (e) Cell-bead co-encapsulation statistics. Abbreviations: C for cell, B for bead, and m for multiple. (f) Comparison of cell capture rate between the present work and several representative reports on single-cell sequencing. (g) Transcript profiling statistics of a Drop-seq run in which HRBs were used.

Encouraged by these findings, we stepped forward to examine bead-cell co-encapsulation (Fig. 4c and Supporting Movie S2). A suspension of HEK293T cells (∼ 200/µL) was co-flowed with the HRBs in the device (Fig. 4c inset micrograph) and encapsulated in the droplets. First, we concluded from the micrograph of Fig. 4d that the overall bead encapsulation rate was 64.1% (n = 136). This is slightly lower than that of our previous results owing to the wider size distribution of Drop-seq bead-based HRBs, but still outperforms Poisson loading (Fig. S5). With respect to the cells, our statistics show that among all the cell-containing droplets, 75% of them contain exactly one bead and one cell (Fig. 4e). To further test the reproducibility, we conducted three repetitive experiments and obtained a total cell capture rate (the ratios of cells being paired with a bead) of 68.8% (n = 3, totally 420 droplets counted), which is over 6-fold higher than the original Drop-seq technique^26^ and surpasses two recent works leveraging on inertial microfluidics^33, 36^ and comparable to a commercial platform that uses soft hydrogel barcoding beads^29^ (Fig. 4f), while maintaining a low rate of doublets, 1.6%.

Having verified that our strategy can significantly improve the bead-cell co-encapsulation, we asked whether the hydrogel coating could potentially leave any side effects. One suspicion was whether the hydrogel coating could be dissolved to expose the functional molecules on the bead surface. To figure it out, we first conducted a preliminary experiment, verifying that the agarose gel coating could be melted by simply heating the droplets at 60 °C without affecting the droplet stability (Fig. S6). Of note, the heating step is also required in the process of reverse transcription, typically conducted at 55-60 °C ^43^. We further asked whether there could be possible residual agarose molecules adsorped on the bead surface, interfering with the subsequent biochemical events. In the case of Drop-seq beads, they are functionalized with single-stranded DNA (ssDNA) oligos to capture the RNA transcripts. Thus, to answer the question, we co-encapsulated them with cells, melted the hydrogel for RNA capturing and conducted a complete library preparation run of Drop-seq. The sequencing data of the prepared cDNA library exhibited a reasonably good quality (Fig. 4g). The number of detected genes and total number of transcripts per barcode group, and the ratio of mitochondria RNA are all comparable to other Drop-seq applications ^33, 36^. These results sufficiently verify that our strategy improves the bead loading and the cell capture efficiency of single-cell transcriptomic sequencing, while not affecting the RNA capturing performance.

## Discussion

As the outer hydrogel of HRBs serves as a physical buffering and lubricating layer, the exact chemical composition of the gel is not critical. Therefore, various kinds of hydrogels can be used to form the HRB’s outer layer^44, 45^. For example, if the current heating procedure to melt the agarose gel is not desired, one could switch agarose to covalently formed and degradable hydrogels such as Polyacrylamide (PAAM)^46^ and poly(ethylene glycol) (PEG)^47^, such that the gel layer can be dissolved chemically or photoresponsively^48^. If necessary, the hydrogel layer can be further modified with probe molecules such as cell-binding peptides^49^ and aptamers^50^ to render the beads additional functionalities. Moreover, the strategy is not dependent on the size and material of the raw beads, as long as the HRBs can be fabricated with the proposed methods. Thus, we expect that our strategy is universally applicable. It should be compatible with various bead materials such as plastics, silica, and ceramics, and compatible with bead diameters ranging from 10 to 100 µm.

Although a simple flow-focusing design was used to load the HRBs into the droplets in the present work (Fig. S2), the HRBs should be readily compatible with various common bead-loading designs, especially those for loading hydrogel beads. In an experiment, we used a tilted-channel design identical to the InDrop platform and obtained a reasonable loading efficiency of ∼50% (Fig. S7). Other than the demonstrated usefulness of improving bead loading efficiency, the combination of a rigid microbead and hydrogel in a core-shell configuration could potentially render bead-based emulsion assays more convenient and microfluidic-free. For rigid microbeads alone, encapsulating them in droplets must rely on microfluidic devices. Herein, the point is that the droplet has to be appropriately larger than the bead to house additional reaction medium and biological targets such as cells. Further, the droplets must be monodispersed to ensure that each reaction is volumetrically identical. Under these requirements, typical microfluidic-free emulsification methods such as vortexing are incapable. However, the HRBs may provide a feasible solution. Uniform HRBs can be pre-manufactured and stored. Once needed, they can be emulsified by vortexing with ease^41^, leading to monodispersed bead-templated emulsions. Afterwards, if necessary, the hydrogel layer can be selectively dissolved for the assays to take place. We anticipate that this strategy should be interesting to future microfluidic-free digital molecular/cellular assays^41, 51, 52^.

## Conclusion

We have developed a general strategy for high-efficiency single-bead encapsulation in microdroplets by using hydrogel coating-assisted close-packed ordering. Using the strategy, the single-bead loading efficiency has exceeded 80% in the demonstrated experiments, which surpasses the Poisson loading and existing inertial ordering-based techniques. Furthermore, we have also shown that using HRBs can substantially increase the cell-bead pairing efficiency in a single-cell sequencing run. The HRBs have been demonstrated to be convenient to manufacture, and the process can be highly scalable. Moreover, the HRBs are comparable with existing microfluidic designs to achieve high-efficiency loading with no requirement for major microfluidic innovations or stringent flow conditions. All of these features show that the strategy is user-friendly to general biological laboratories. Thus, we anticipate that the strategy holds the potential to be broadly applied for high-efficiency single-cell analysis and digital assays. Our current efforts are being paid to develop microfluidic-free single-cell assays with these HRBs.

## Author contributions

Conceptualization: YL, JZ, XM; methodology: YL, LC; investigation: LC, YZ, JL, CX, YX, CT, RZ; visualization: LC, YL; supervision: YL, JZ, XM; writing – original draft: YL, LC; writing – review & editing: YL, LC.

## Acknowledgements

This work is supported by the National Natural Science Foundation of China (Grant No. 61904105) and the start-up funding of ShanghaiTech University. The authors also acknowledge the sponsorship from Double First-Class Initiative Fund of ShanghaiTech University. YL would like to thank the support from Shanghai Clinical Research and Trial Center. The fabrication of the microfluidic devices was partially carried out in the Soft Matter Nanofabrication Laboratory of SPST.

