## Supporting_Information for "Exceeding 80% efficiency of single-bead encapsulation in microdroplets through hydrogel coating-assisted close-packed ordering": Supporting_Information_V2.pdf

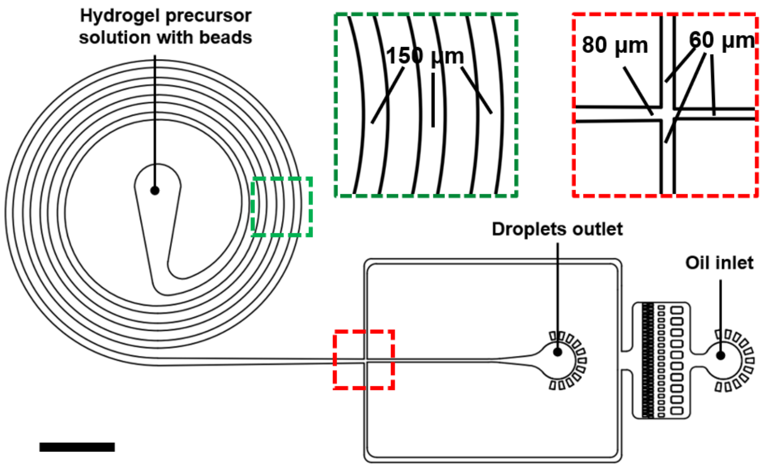


**Figure S1.** The layout of a jetting microfluidics device for manufacturing the hydrogel-coated rigid beads (HRBs). The insets highlight the geometry of the curve channel (green dash lines) and the flow-focusing junction (red dash lines). The scale bar denotes 1.5 mm.

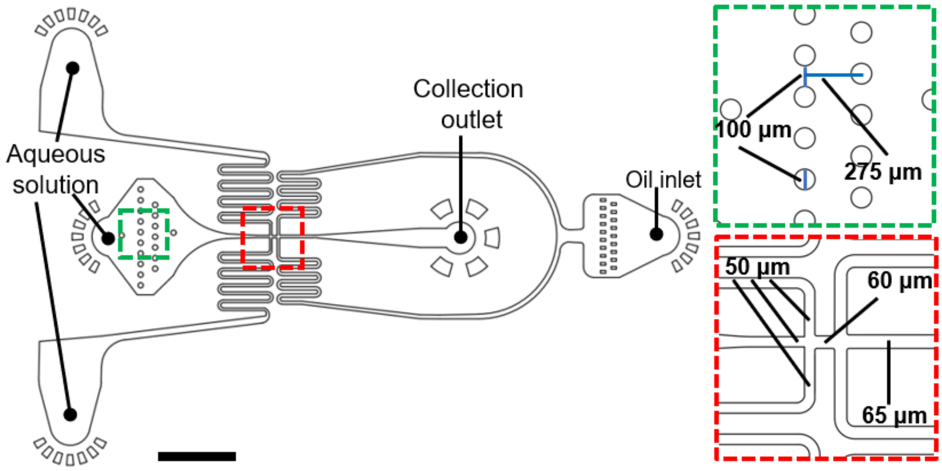


**Figure S2.** The layout of a dual-junction flow-focusing device for loading the HRBs into droplets. The insets detail the geometry of the pillar filter (green dash lines) and the junctions (red dash lines). The scale bar denotes 1.5 mm.

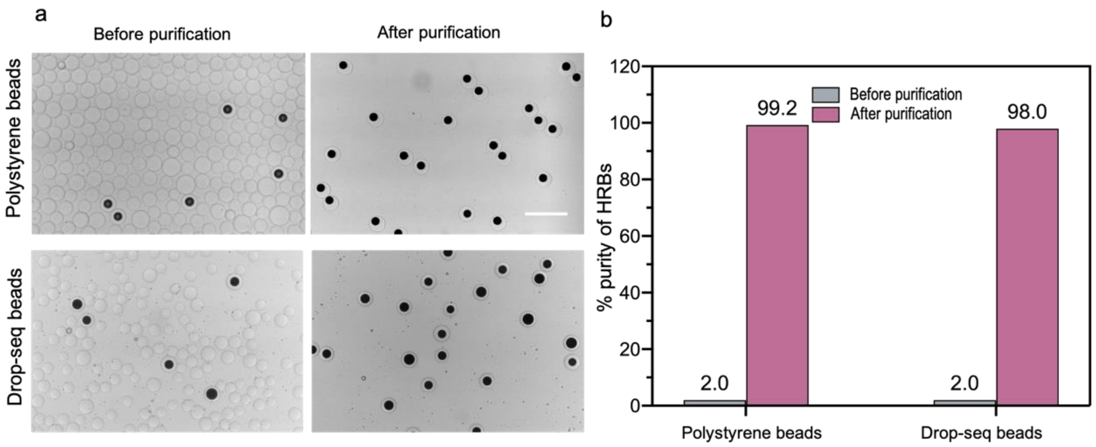


**Figure S3.** Purification of HRBs. (a) micrographs of HRBs right after manufacturing and after a centrifugal purification process. Scale bar: 150 µm. (b) Statistics of HRBs purity before and after purification (n = 323 for polystyrene beads and 345 for Drop-seq beads).

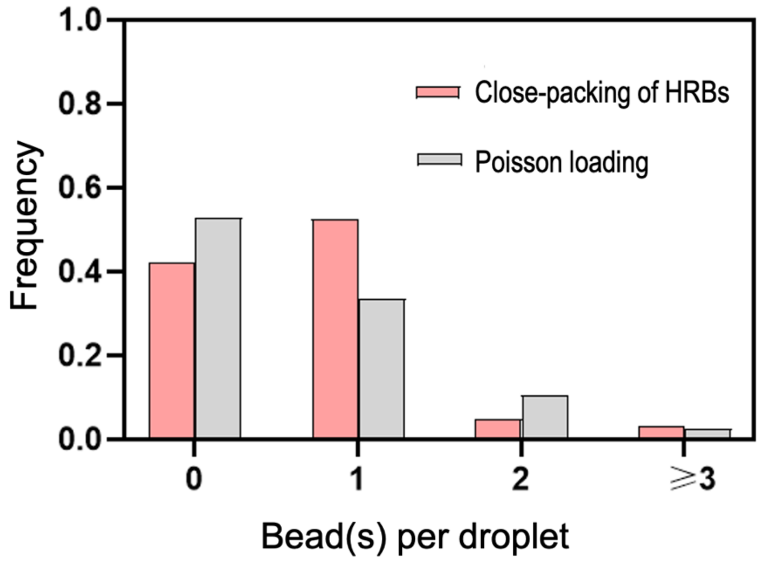


**Figure S4.** The comparision between close-packed loading of vortex-made HRBs and Poisson loading ($\lambda$= 0.634).

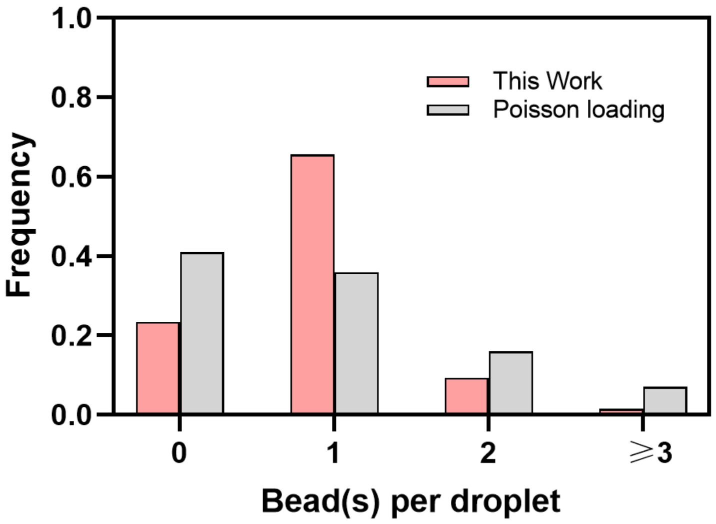


**Figure S5.** Comparison of loading Drop-seq beads using the HRB stratey and Poisson loading at the same bead density (λ= 0.89).

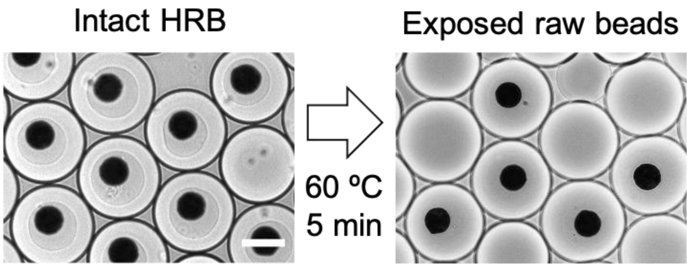


**Figure S6.** Dissolution of the agarose hydrogel coating of HRBs by heating to expose the raw beads. Scale bar: 50 μm.

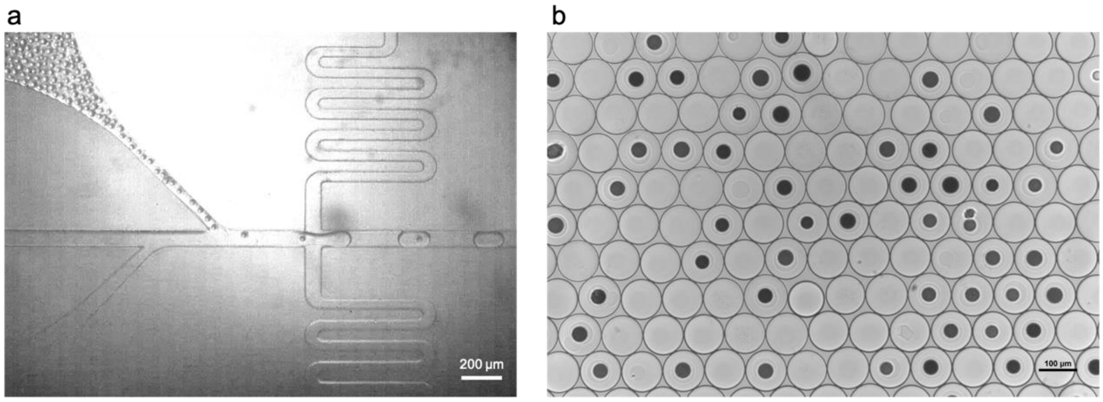


**Figure S7.** (a) Loading Drop-seq bead-based HRBs using a titled channel microfluidic design and (b) the generated droplets featuring a bead loading efficiency of 48.4% (n > 300).
